# Spatial control of myosin regulatory light chain phosphorylation modulates cardiac thick filament mechano-sensing

**DOI:** 10.1101/2025.04.25.650613

**Authors:** Caterina Squarci, Daniel Koch, Paul Aanaya, Kenneth S. Campbell, Thomas Kampourakis

**Affiliations:** Division of Cardiovascular Medicine, Internal Medicine, College of Medicine, University of Kentucky, Lexington KY 40536, USA; Max Planck Institute for Neurobiology of Behavior — CAESAR, Ludwig-Erhard-Allee 2, D-53175 Bonn; Randall Centre for Cell and Molecular Biophysics, King’s College London, London SE1 1UL, United Kingdom

**Keywords:** cardiac muscle regulation, cardiac myosin, phosphorylation

## Abstract

The heart can adapt its performance in response to changing metabolic demands of the rest of the body. A central mechanism intrinsic to the heart is to modulate the function of the cardiac contractile proteins via post-translational modifications. Although phosphorylation of the cardiac myosin motor-associated regulatory light chain (RLC) by cardiac myosin light chain kinase (cMLCK) has been recognized as a key signalling pathway to increase myocardial contractile function, little is known about its molecular mechanism of action. Here, we show that phosphorylation of RLC is not a stochastic process but a spatially tightly controlled mechanism. Myosin motors in the region of the thick filament associated with cardiac myosin binding protein-C (cMyBP-C) are the primary target for phosphorylation by cMLCK. Moreover, phosphorylation of RLC likely only leads to activation of one of the two myosin motors of the cardiac myosin molecule and increases their force-dependent recruitment. We propose that RLC phosphorylation exerts its functional effects via increasing the gain of the mechano-signalling between different zones of the thick filament. A better mechanistic understanding of the role of RLC phosphorylation likely underpins the development of therapeutic interventions for both heart disease and heart failure.

## Introduction

The cardiac contraction-relaxation cycle is driven by the cyclic interactions of the actin-containing thin and myosin-containing thick filaments organized into the highly ordered sarcomeres (1). Calcium binding to the troponin complex in the thin filaments allows tropomyosin to azimuthally rotate around the thin filament and expose myosin-binding sites on actin. Subsequently, the myosin motor domain (or head) from the neighbouring thick filaments can attach to actin. Through hydrolysis of ATP, the formed actin-myosin complex undergoes the working stroke, which drives cardiac muscle force development and shortening. Conversely, calcium release from troponin switches the thin filaments OFF followed by detachment of myosin motors from actin and mechanical relaxation of the heart. In this actin-centric view of regulation, contraction is solely controlled by the calcium-dependent structural changes in the thin filaments that determine the availability of myosin-binding sites on actin.

However, myosin-based regulation of cardiac contractile function has emerged as new fundamental concept with important implications for both the physiological and pathophysiological control of heart muscle performance (2, 3). Although originally identified in invertebrate muscles, the concept of a regulatory structural change in the myosin-containing thick filament was later also applied to mammalian striated muscles, including the thick filaments of heart muscle (4-9). According to this concept, the cardiac thick filaments transition from a diastolic OFF state, characterized by myosin heads adopting an asymmetrical folded state or interacting heads motif (IHM) in a quasi-helical arrangement on the surface of the thick filament backbone, to a systolic ON state that allows actin binding and force generation. The myosin motor OFF state is stabilized by intermolecular interactions between the two myosin heads of the dimeric myosin molecule and their tail domains, and intramolecular interactions between myosin heads on adjacent crowns and other thick filament components such as cardiac myosin binding protein-C (cMyBP-C) and titin (8, 9). The rate of transition between those states likely controls the rate of isovolumetric contraction and relaxation, and the duration of the ejection phase (10-12).

Phosphorylation of the myosin motor domain-associated regulatory light chain (RLC) is an important determinant of cardiac muscle function. Ablation of RLC phosphorylation leads to heart disease and heart failure in transgenic animal models. In contrast, increasing RLC phosphorylation is cardioprotective and increases the performance of the failing heart muscle (13-16). Moreover, modulation of RLC phosphorylation has been proposed as a potential therapeutic intervention for heart disease and heart failure (16, 17).

The level of RLC phosphorylation is physiologically controlled by the activity of the cardiac isoform of myosin light kinase (cMLCK), and the myosin phosphatase complex comprised of the catalytic subunit of protein phosphatase-1 (PP1) and a regulatory subunit (myosin phosphatase targeting subunit-2, MYPT2) (18-20). cMLCK activity is primarily regulated via its calcium and calmodulin-dependent activity, whereas the regulation of the myosin phosphatase is less well understood (21).

RLC phosphorylation increases the force production, calcium sensitivity and crossbridge kinetics of isolated cardiac muscle, as well as an increased stretch-activation response, which translates into an increased cardiac output on the organ level (22-25). Moreover, a gradient of RLC phosphorylation through the ventricular wall and from the base to the apex was proposed to increase the torsional movement of the heart during systole and facilitate ejection (25, 26).

Many previous studies have attempted to elucidate the underlying mechanistic basis for the functional effects of RLC phosphorylation. Early studies on isolated invertebrate and vertebrate skeletal thick filaments suggested that RLC phosphorylation destabilizes the folded OFF state of the myosin motors (27, 28), although this mechanism has not been generally accepted (11, 24). Other potential candidate mechanisms are an increase in force production per myosin motor and signalling between thick and thin filaments (24, 25, 29).

In the present study, we explore the molecular mechanism of RLC phosphorylation using a wide range of biochemical, biophysical and super-resolution imaging techniques. The results presented below show that (I) RLC phosphorylation is not a stochastic process but more likely a tightly controlled mechanism in the sarcomere. (ii) RLC phosphorylation by cMLCK is largely restricted to the inner two-thirds of the thick filaments, corresponding to the region of the filament associated with cMyBP-C. Moreover, using fluorescence polarization measurements in comparison with recently published high-resolution structures of the isolated cardiac IHM, we show that (iii) RLC phosphorylation likely only affects the conformation of one of the two myosin heads of the double-headed myosin molecule and increases their force-dependent recruitment. Our results can be summarized in a new model suggesting that RLC phosphorylation exerts its functional effects by increasing the gain of the mechano-signalling between different zones of the thick filament.

## Results

### Heterogenous phosphorylation of myosin heads in intact thick filaments by cMLCK

Zonal differences in the thick filament structure (e.g. D-vs C-zone) associated with different titin super-repeats and accessory proteins (e.g. cardiac myosin binding protein-C, cMyBP-C) and the asymmetric OFF conformation of the double-headed cardiac myosin molecule (called the ‘interacting heads motif’, IHM; **Fig. 1a**) suggest that multiple conformations of myosin heads co-exist in the native thick filament (8, 9, 30). We hypothesized that these different populations of myosin heads might be differentially phosphorylated by the cardiac isoform of myosin light chain kinase (cMLCK).

**Figure 1.**
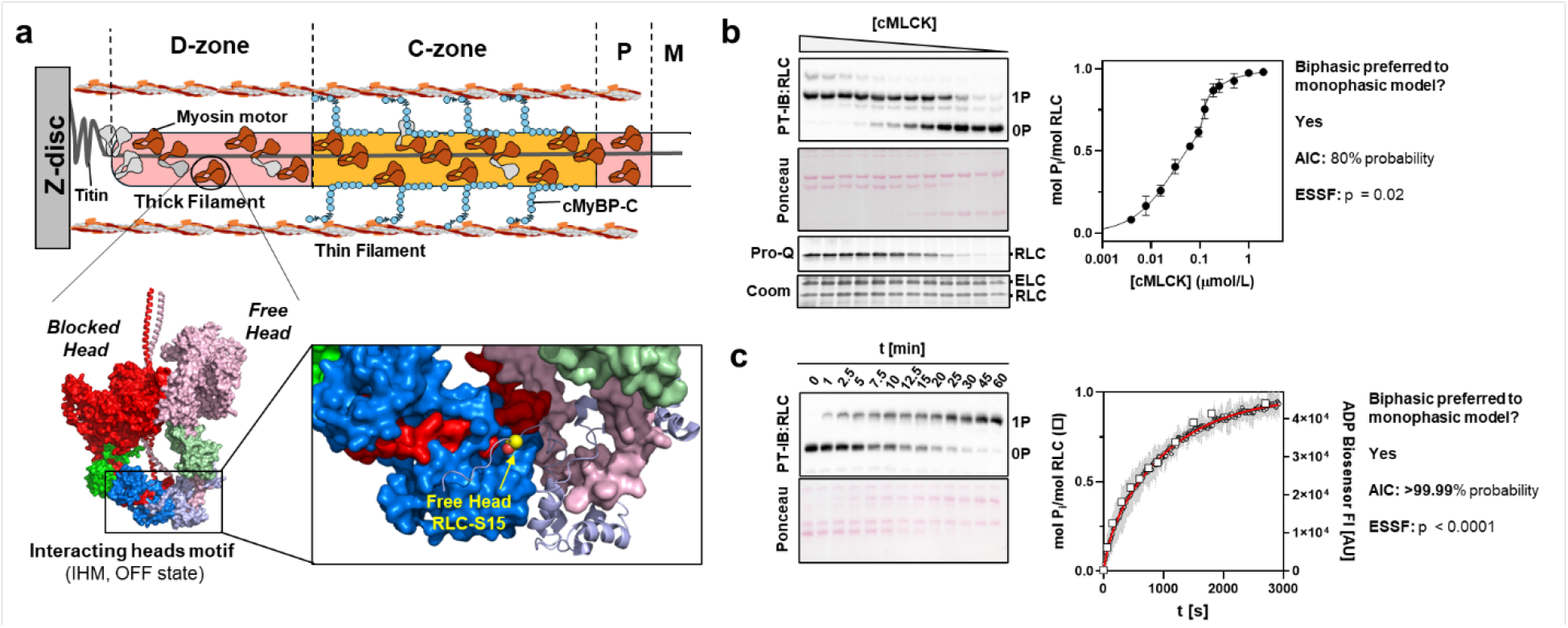
Non-uniform phosphorylation of RLCs by cMLCK in intact thick filaments. (a) Top: Cartoon representation of the half-thick filament with D-, C- and P-zone, and M-band labelled accordingly. Bottom: Atomic model of the isolated human cardiac interacting heads motif (IHM). The free and blocked head are labelled accordingly. The insert on the right shows a magnified view of the RLC-region of the IHM with the phosphorylatable RLC residue highlighted in yellow. (b) Steady-state dose-response relation of RLC phosphorylation in isolated ventricular myofibrils in the presence of fixed concentration of blebbistatin and protein phosphatase 1, and increasing concentrations of activated cardiac myosin light chain kinase (cMLCK). Phosphorylation levels were analyzed by Phostag™-Western blot against RLC (top), separating unphosphorylated (0P) and phosphorylated RLC (1P) for n=3 independent myofibril preparations, and ProQ Diamond staining (bottom). Continuous line shows biphasic fit to data points. (c) Time-dependent phosphorylation of RLC in relaxed ventricular myofibrils analyzed by Phostag™-Western blot against RLC (open squares, n=1) and ADP-biosensor assay (open circles, grey area denotes 95% CI, n = 3 independent myofibril preparations). The continuous red line indicates bi-exponential fit to data points.

To test this idea, we incubated cardiac myofibrils isolated from rat ventricular tissue, which contains both intact thick and thin filaments organized in the native myofilament lattice, with recombinant cMLCK in the presence of ATP, calmodulin (CaM) and Ca^2+^. We have previously shown that the basal RLC phosphorylation in these samples is less than 0.05 mol P_i_ mol RLC^-1^, and that exogenous cMLCK specifically phosphorylates RLC under these conditions (18, 24) (**Fig. S1a**). To stabilize the OFF state of the myosin motors and prevent contraction of myofibrils during the kinase assay, we tested two inhibitors of cardiac myosin, butadiene-monoxime (BDM) and Blebbistatin and analysed the time-dependent phosphorylation of RLC using Phostag^TM^-Western-blot (**Fig. S1b**). Although cMLCK phosphorylated RLCs in cardiomyofibrils (CMFs)with about the same kinetics in the presence of either BDM or Blebbistatin (k_app_ of about 0.06 mol P_i_ min^-1^), the maximal level of phosphorylation was significantly lower in the presence of BDM (65% for BDM vs 95% for Blebbistatin). It was previously suggested that BDM might acts as a chemical phosphatase for RLC (31). We tested this idea by incubating isolated fully phosphorylated RLC either in the absence or in the presence of 25 mmol L^-1^ BDM at 25ºC. BDM treatment did not lead to any significant dephosphorylation of RLC within three days, suggesting that BDM does not act as a chemical phosphatase. Similarly, Blebbistatin and BDM had no effect on the activity of cMLCK towards isolated RLC (**Fig. S1e**). We conclude that the lower level of RLC phosphorylation in the presence of BDM is likely associated with the different biochemical or structural states of the cardiac myosin motors in the presence of the two compounds.

Next, we incubated isolated rat cardiac myofibrils in the presence of 25 μmol L^-1^ Blebbistatin with increasing concentrations of CaM/Ca^2+^-activated cMLCK. We also added a fixed concentration of recombinant protein phosphatase-1 (PP1, 0.1 μmol L^-1^) to the myofibrils to allow dephosphorylation of RLCs and incubated the mixtures for 90-120 mins at 30ºC to reach a steady state level (**Fig. 1b**). Analysis by Phostag^TM^-SDS-PAGE and Western-blot against RLC showed a dose-response curve that is best described by a biphasic relationship between the level of RLC phosphorylation and cMLCK concentration with EC_50_ of about 0.01 and 0.1 μmol L^-1^. This suggests that myosin heads in intact myofilament lattice are not uniformly phosphorylated by cMLCK and that least two populations of myosin heads co-exist in the myofibrils that are phosphorylated by cMLCK with significantly different efficacies.

To further test the hypothesis of multiple populations of myosin heads, we next determined the kinetics of RLC phosphorylation by cMLCK in myofibrils (**Fig. 1c**). Initial experiments were analyzed using Phostag^TM^-Western-blot (**Fig. 1c**, open squares). However, the technique has an effective time resolution in the minutes time-scale, which might be too slow to accurately capture the phosphorylation kinetics. We therefore utilized a methodology with a higher time resolution based on a fluorescent ADP biosensor (32) (**Fig. 1c**, open circles). ADP is the other byproduct of the phosphorylation reaction and linearly related to the amount of phosphate transferred from ATP to RLC via cMLCK. Both methodologies are in very good agreement with each other and show that the RLC phosphorylation kinetics are best described by a bi-exponential process with two rate constants that differ by about one order of magnitude. This is in excellent agreement with the steady-state data shown above, supporting the idea of multiple population of myosin heads that are differentially phosphorylated by cMLCK.

As a control we measured the time-dependent phosphorylation of isolated recombinant RLC via both Phostag^TM^-SDS-PAGE and ADP biosensor assay. As expected, the results from both assays were best fitted by a monophasic model (i.e. single exponential) (**Fig. S2a**), suggesting a kinetically homogenous first-order reaction. We also used a construct composed of two RLCs bound the proximal region of myosin S2, called miniHMM, as a substrate in kinase assays (**Fig. S2b**) (33). However, strikingly, the phosphorylation reaction as reported by changes in the ADP biosensor follows a bi-exponential kinetic, suggesting that the two phases observed in the myofibril experiments might be at least partly associated with the two heads of the asymmetric myosin molecule.

### cMLCK predominantly phosphorylates myosin heads in the thick filament C-zone

The results presented above suggest the existence of at least two different populations of myosin heads in intact myofibrils that are phosphorylated with different efficacies by cMLCK. Next, we used an antibody against phosphorylated RLC (RLC-Ser15P, green) in combination with super-resolution microscopy to determine the spatial distribution of RLC phosphorylation along the thick filaments in the ventricular myofibril samples (**Fig. 2a**). Myofibrils were counter-stained using an antibody raised against the N-terminal region of the myosin heavy chain (MHC, red) to label all myosin heads independently of their phosphorylation level. Specificity of the anti-RLC-Ser15P antibody was validated by both Western-blot and immuno-staining (**Fig. S3**) (34).

**Figure 2.**
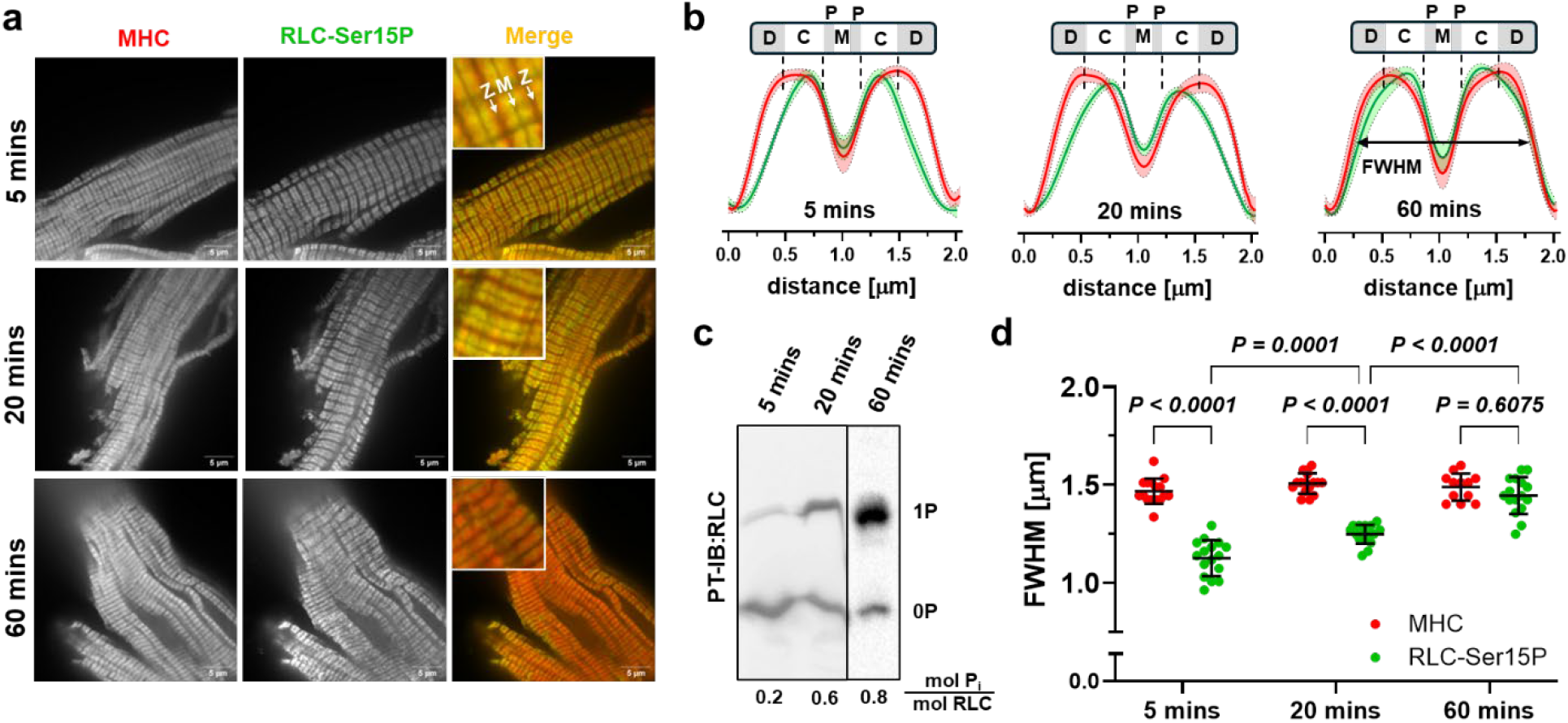
cMLCK preferentially phosphorylates myosin heads in the C-zone. (a) Super-resolution imaging of rat ventricular myofibrils stained against myosin heavy chain (MHC, red) and phosphorylated RLC (RLC Ser15P, green) at different time points after incubation with activated cMLCK. (b) Normalized averaged intensity profiles (MHC, red; RLC-Ser15P, green) after 5 mins, 20 mins and 60 mins of incubation of cardiac myofibrils with activated cMLCK with shaded areas denoting 95% confidence intervals. The position of the thick filament D-, C-, P-zones and M-band are shown to scale above the plots. The full width half maximum (FWHM) is indicated by a double arrow. (c) Phostag™-Westen-blot against RLC of cardiac myofibrillar samples with the average RLC phosphorylation in mol Pi mol RLC^-1^ level shown below. (d) Comparison of the full width half-maximum (FWHM) of the MHC (red) and RLC (green) intensity profiles (n=16 for 5 mins, n=16 for 20 mins and n=16 for 60 mins, for n=3 independent repeats). Statistical significance of differences between values was assessed with a two-way ANOVA followed by Tukey’s multiple comparison test.

The normalized average intensity profiles for both MHC (red) and RLC-Ser15P (green) are shown in **Figure 2b**. Strikingly, 5 mins and 20 mins of incubation of myofibrils with cMLCK, corresponding to intermediate levels of RLC phosphorylation with about 0.2 and 0.6 mol P_i_ mol RLC^-1^ (**Fig. 2c**), respectively, resulted in phosphorylation of myosin heads predominantly in the inner two-thirds of the thick filament. In contrast, myosin heads towards the tip of the thick filaments (towards the Z-line) were less or not phosphorylated under these conditions and only prolonged incubation (60 mins, >0.8 mol P_i_ mol RLC^-1^) led to a more homogenous RLC phosphorylation along the whole length of the thick filaments. We quantified the spatial distribution of RLC phosphorylation along the thick filaments by comparing the full width half maximum (FWHM) of the MHC and RLC-Ser15P intensity profiles. The FWHM of the MHC profile was about 1.5 μm under all conditions tested, which is in agreement with the estimated length of the A-band of ventricular myofibrils of 1.52-1.58 μm as measured by electron microscopy (35, 36). In contrast, after 5 mins and 20 mins of incubation, the RLC phosphorylation showed a significantly smaller FWHM than the MHC signal, whereas after 60 mins the differences was no longer statistically significant (**Fig. 2d**). The average FWHM of the RLC-Ser15P signal after 5 mins of cMCLK incubation (about 1.1 μm) is in excellent agreement with the doublet width of the thick filament C-zone as estimated by electron microscopy of rat myofibrils and cryoEM reconstructions of mouse thick filaments in intact ventricular trabeculae (both ∼1 μm), and super-resolution microscopy measurements of fluorophore-conjugated cMyBP-C expressed in isolated rat cardiomyocytes (∼1.1 μm) (9, 35, 37). Noteworthy, the FWHM of the RLC-Ser15P profile slightly increases from about 1.1 μm to 1.25 μm when the RLC phosphorylation level increases from 0.2 to 0.6 mol P_i_ mol RLC^-1^, respectively. Since the C-zone contains about 50% of the total number of myosin motors in rodent cardiac thick filaments, the quantitative comparison suggests that all myosin motors in the C-zone are phosphorylated first before myosin motors in the D-zone become phosphorylated.

We also isolated myofibrils from human donor myocardium under native conditions with an RLC phosphorylation level of about 0.5 mol P_i_ mol RLC^-1^, which is in good agreement with previous reports on intact cardiac preparations (34). Similar to the results obtained by in-vitro phosphorylation of rat cardiac myofibrils, staining against both Ser15-phosphorylated RLC and MHC showed that myosin motors are preferentially phosphorylated in the C-zone in human myofibrils under basal conditions (**Fig. S4**).

### RLC phosphorylation activates the blocked head of cardiac myosin

In order to gain more insights into the structural effects of RLC phosphorylation on the myosin head conformation in intact thick filaments, we compared the previously published orientation distributions of the RLC-region of cardiac myosin based on fluorescent polarization measurements in demembranated rat ventricular trabeculae with recent high-resolution cryo-electron microscopy (cryoEM) structures of the isolated human cardiac IHM (30, 38) (**Fig. 3a**).

**Figure 3.**
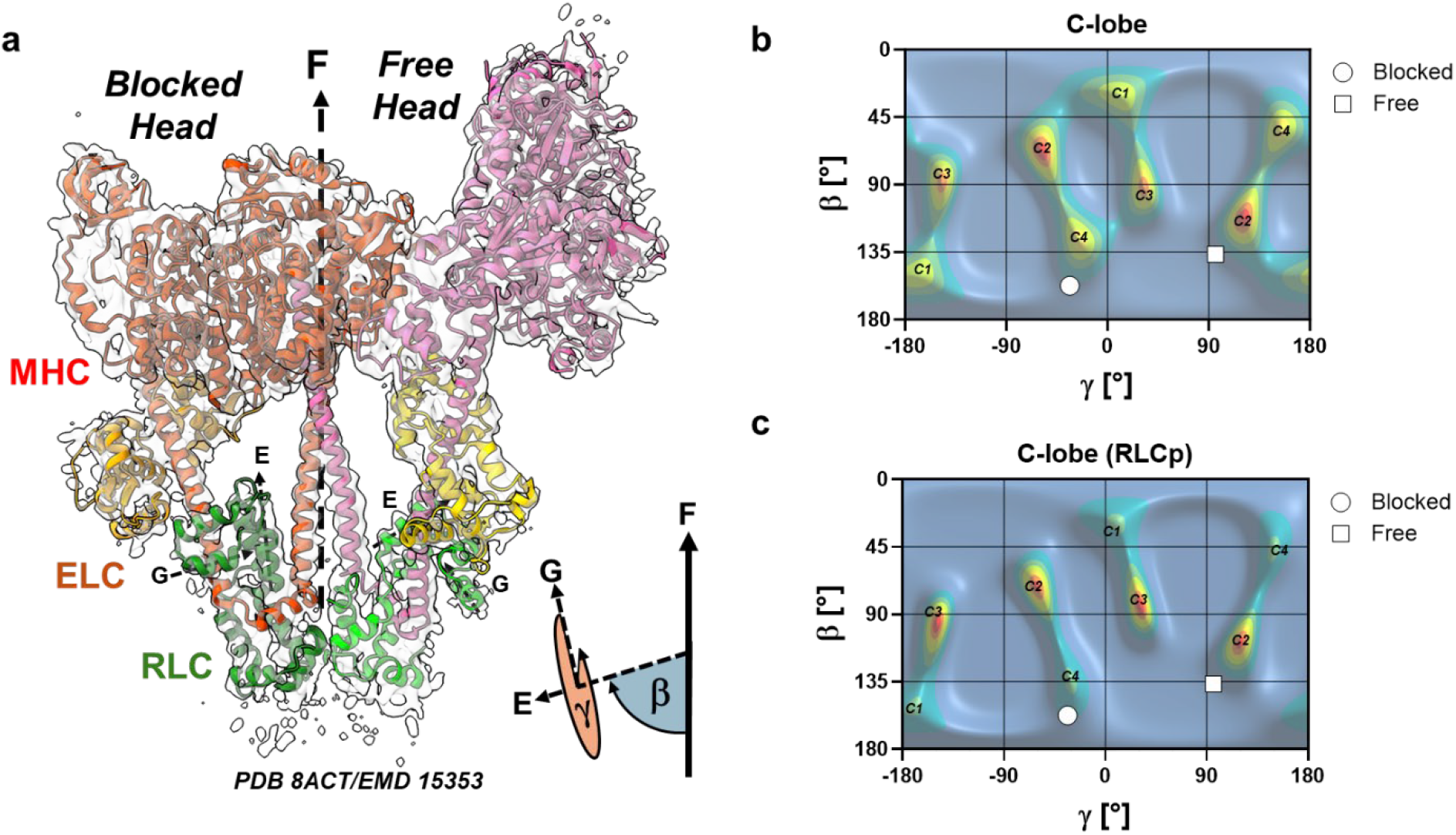
RLC phosphorylation activates the blocked head. (a) Atomic model (PDB 8ACT) of the isolated human cardiac interacting heads determined by cryo-electron microscopy (EMD 15353). The electron density map (transparent surface) is shown superimposed to the atomic model of the IHM shown in cartoon representation (MHC-myosin heavy chain, ELC – essential light chain, RLC – regulatory light chain). The E-and G-helix of the RLCs are labelled accordingly. The cartoon at the bottom right shows the definition of the RLC orientation determined by the Euler angles β and γ. F denotes the orientation of the filament long axis. (b) Maximum entropy (ME) distribution of the RLC C-lobe orientation before RLC phosphorylation using the atomic coordinates from PDB 8ACT. (c) Maximum entropy (ME) distribution of the RLC C-lobe orientation during full calcium activation after RLC phosphorylation using the atomic coordinates from PDB 8ACT. The calculated orientations of the RLC of the blocked and free head based in the structure of the isolated IHM (PDB 8ACT) are indicated by white circles and squares, respectively.

The orientations of the RLC-region determined by polarized fluorescence are constrained by the two Euler angles β and γ, with β corresponding to the angle between the RLC E-helix and the filament axis, and γ describing the rotation of the RLC around the E-helix vector (**Fig. 3a**). The orientation distribution of the RLC in the unphosphorylated state was determined by polarized fluorescence from a bifunctional rhodamine probe attached to four sites on the RLC and analyzed using a maximum entropy (ME) formalism (18, 39) (**Fig. 3b**). The relaxed ME map is dominated by four peaks, which were designated as C1 through C4. Since polarized fluorescence cannot distinguish between solutions of (β,γ) and (180-β,γ+180), every population appears twice in the ME maps and can be considered as solutions corresponding to the two sides of the bipolar thick filaments.

The calculated (β,γ) orientations of the RLC-region of the so-called free- and blocked-head from the isolated human IHM are in very good agreement (within the experimental error of the techniques (10º-20º)), with orientations C2 and C4 in the relaxed ME maps, respectively (**Fig. 3b**). This suggests that that peaks C2 and C4 likely correspond to myosin heads adopting free and blocked head orientations of the IHM state in the relaxed ventricular trabeculae. However, the presence of additional peaks (C1 and C3) indicates that not all myosin motors adopt this folded OFF state under these conditions.

Phosphorylation has significant effects on the RLC orientation distribution during the relaxed state (**Fig. c**). Although the position and intensity of peak C2 did not change significantly, peak C3 become more dominant. More strikingly, however, both peaks C4 and C1 almost completely disappeared, suggesting that the blocked head of the IHM changes its conformation after RLC phosphorylation and that likely the free head remains in the folded OFF state.

Taken together, the comparison of the fluorescence polarization data from intact myofilaments with the cryoEM reconstructions of the isolated IHM suggests that RLC phosphorylation primarily affects the conformation of the so-called blocked head RLC of the asymmetric myosin molecule and that the free head RLC likely remains in the IHM state.

### Spatially explicit modelling of the functional effects of RLC phosphorylation is consistent with activation of a single head in the C-zone

Phosphorylation of RLC has been shown to modulate cardiac muscle force production in a sarcomere length-dependent manner (24, 40) (**Fig. 4a**). At short sarcomere length (∼1.9 μm) RLC phosphorylation increases the calcium sensitivity of force production as measured by an increase in pCa_50_ by about 0.08 (pCa = -log_10_[Ca^2+^] for half-maximal activation) from 5.56 to 5.64 but had no effect on the steepness/cooperativity of the force-pCa relation. In contrast, RLC phosphorylation at long sarcomere length (∼2.3 μm) increased the calcium sensitivity of force by about 0.12 pCa and significantly decreased its cooperativity, leading to a large increase in isometric force production in the physiological range of calcium concentrations close to pCa 5.8.

**Figure 4.**
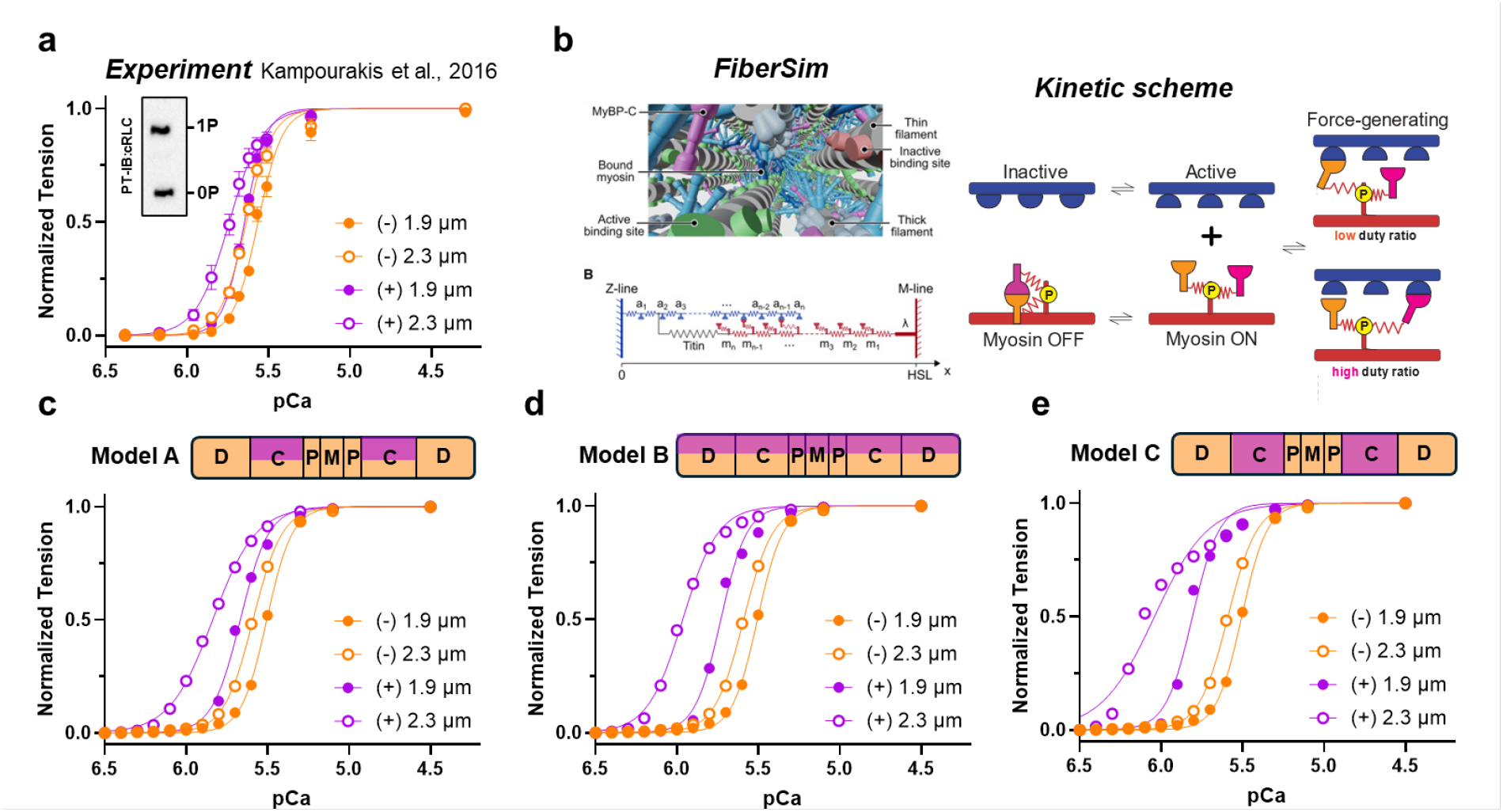
Spatially-explicit modelling is consistent with activation of the blocked head in the C-zone. (a) Force-pCa relation of ventricular trabeculae before (-) and after RLC phosphorylation by cMLCK (+) at short (1.9 μm) and long (2.3 μm) sarcomere length (n=5 trabeculae for each condition). The inset shows the RLC phosphorylation level of trabeculae after cMLCK treatment. Data from Kampourakis et al., 2016 (24). (b) FiberSim core model (left) and kinetic scheme (right). Simulated force-pCa relations for (c) activation of the blocked by RLC phosphorylation only in the C-zone (model A), (d) activation of the blocked head in the whole thick filament (Model B) and (e) activation of both blocked and free head in the c-zone (Model C).

We used a spatially-explicit model of the sarcomere incorporating a force-dependent recruitment of myosin motors from the OFF state (i.e. FiberSim (41)) to test if the spatial distribution and structural effects of RLC phosphorylation described above are consistent with the experimental data (**Fig. 3b**). The model uses a kinetic scheme whereby RLC phosphorylation increases the force-sensitive duty ratio of only one of the two heads of a dimeric myosin molecule, corresponding to the results of the fluorescence polarization experiments (**Fig. 3**) showing that RLC phosphorylation has different effects on the two heads of the dimeric myosin molecule.

In the first model, RLC phosphorylation was restricted to the C-zone of the thick filament (**Fig. 4c**, Model A), which is consistent with the imaging data shown in **Figure 2**. The simulated force-pCa curves at both sarcomere lengths are qualitatively in very good agreement with the experimental data. Either phosphorylation or increase in sarcomere length increases calcium sensitivity by about the same magnitude without affecting the steepness of the force-pCa relation, leading to almost identical force production at around physiological relevant [Ca^2+^] (pCa 6-5.8). In contrast, RLC phosphorylation at long sarcomere length increased calcium sensitivity but also decreases the steepness/cooperativity of force production, leading to a large increase in isometric force at sub-optimal [Ca^2+^].

As controls, we also simulated a homogenous RLC phosphorylation distribution along the whole of the thick filament (**Fig. 4d**) and activation of both myosin heads in the C-zone via RLC phosphorylation (**Fig. 4e**) using the same kinetic parameters as in Model A. Both models did not recapitulate the experimental data, supporting the hypothesis that the functional effects of RLC phosphorylation are controlled by both its effect on the asymmetric myosin molecule and its spatial distribution within the thick filament.

### RLC phosphorylation increases the mechano-sensitivity of myosin head activation

Modelling of the experimental data using FiberSim suggested that the primary effect of RLC phosphorylation is to increase the force-dependent recruitment of myosin motors from the folded OFF state (42).

We tested this hypothesis by using a slack-re-stretch protocol of Ca^2+^-activated demembranated rat right ventricular trabeculae. In this protocol trabeculae are mechanically unloaded by an initial step release to allow unloaded shortening, which is followed by a re-stretch to the original length to restore myofilament load and force development to pre-release levels (43).

In addition to the mechanical data, we monitored the conformation of the myosin motors by exchanging the native RLCs in the demembranated trabecula preparations with a recombinant RLC crosslinked to a bifunctional rhodamine probe along its E-helix (10, 38) (**Fig. 5a**). This allows to determine the orientation of the RLC-region of the myosin motors with respect to the filament axis using polarized fluorescence with milli-second time-resolution. The orientation of the probe (and therefore the myosin motors) is described by the order parameter <*P*_*2*_>, which ranges from -0.5 to 1 for the RLC E-helix being either perpendicular or parallel to the filament axis, respectively (44).

**Figure 5.**
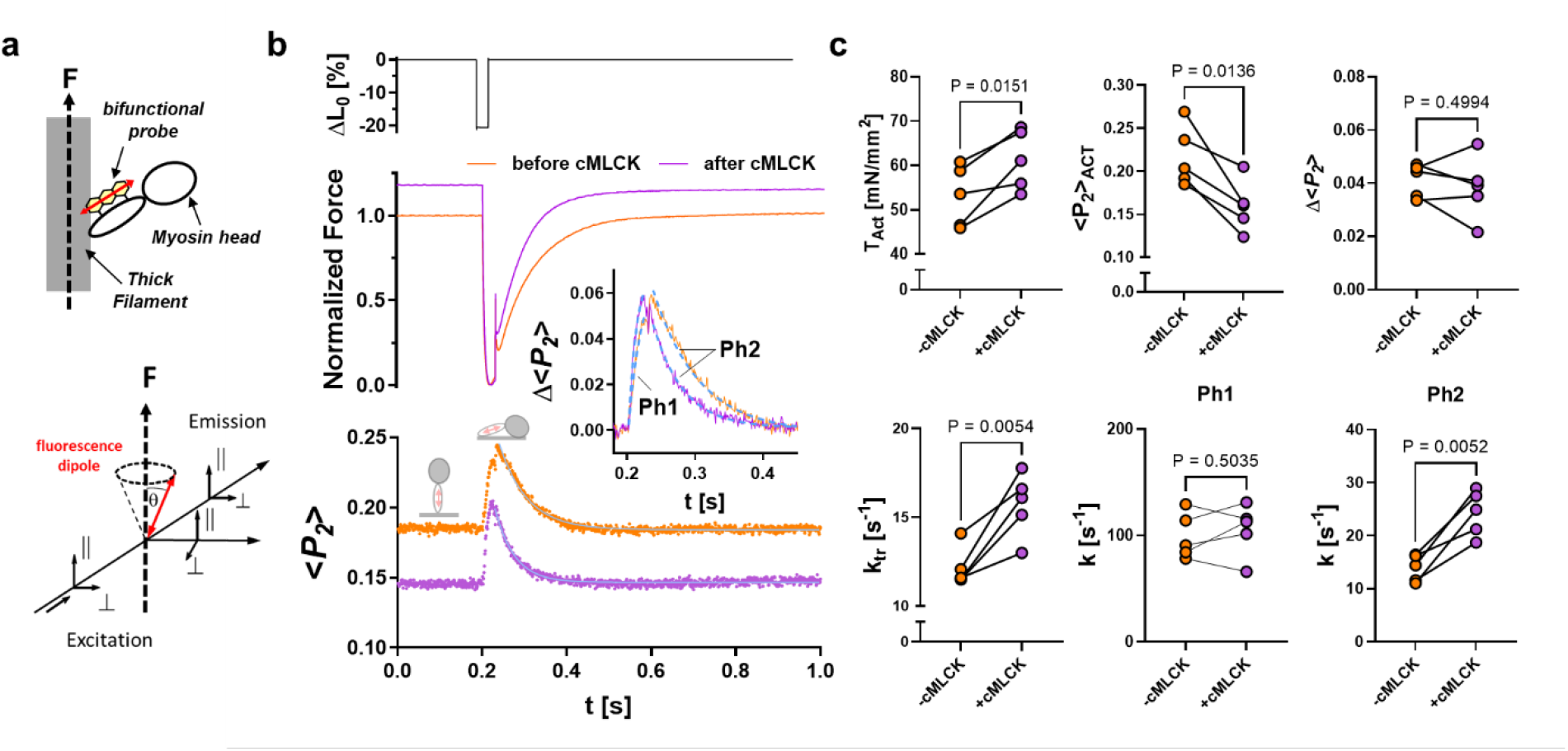
Changes in the orientation of the cRLC E-helix probe in ventricular trabeculae in response to slack-stretch protocol before and after cMLCK treatment. (a) Cartoon of the polarized fluorescence assay to determine the RLC E-helix orientation. (b) Representative traces of muscle length (top), normalized force (middle) and <*P_2_*> (bottom) during the slack-stetch protocol before (orange) and after RLC phosphorylation by cMLCK (purple). (c) Summary of steady state and transient force and <*P_2_*> parameters for n=5 independent trabeculae preparations. Statistical significance of difference between values were assessed with a two-sided, paired student’s t-test.

RLC-exchanged trabeculae were Ca^2+^-activated, and once steady-state force was established, the trabeculae were rapidly shortened and re-stretched. Representative length, force and <*P*_*2*_> traces before and after phosphorylation of RLC by exogenous cMLCK are shown in **Figure 5b**. RLC phosphorylation by cMLCK significantly increases both isometric force at full Ca^2+^-activation and the rate of force re-development (k_tr_) (**Fig. 5c**). In excellent agreement, the value for <*P*_*2*_> during steady-state activation (<*P*_*2*_>_ACT_) significantly decreased after RLC phosphorylation, suggesting that myosin heads are more perpendicular (and therefore more activated).

The transient changes in the myosin head orientation during the release-restretch protocol are best described by two phases. Phase 1 (Ph1) describes the partial recovery of the <*P*_*2*_> towards

the relaxed value directly after the step-release, which is likely associated with detachment of myosin heads from actin and partial recovery of the myosin head OFF state (10). Surprisingly, RLC phosphorylation neither affected the rate of Ph1 nor its amplitude (Δ<*P*_*2*_>), suggesting that RLC phosphorylation does not affect the detachment kinetics of myosin from actin under unloaded conditions nor the rate at which myosin heads reform the folded parallel OFF state.

In contrast, RLC phosphorylation significantly increased the rate of Phase 2 (Ph2) of the <*P*_*2*_> transient directly after the re-stretch, which is associated with myosin motors leaving the folded OFF state, attachment to actin and force-generation. Interestingly, however, although the rate of k_tr_ and Ph2 are almost identical before RLC phosphorylation (rates of ∼13s^-1^ for both), the rate of Ph2 (∼24 s^-1^) is significantly faster than k_tr_ (∼16 s^-1^) after RLC phosphorylation. This can be explained by a faster force-dependent transition of the myosin motors from the OFF to the ON state after RLC phosphorylation that precedes force generation.

Taken together, these results indicate that RLC phosphorylation increases force-dependent activation of the myosin motors, i.e. it increases the gain of the mechano-sensing mechanism of the myosin filament. We further tested this hypothesis by repeating the experiment but omitting the re-stretch so that the thick filaments remain mechanically unloaded (**Fig. S5**). Although RLC phosphorylation also increased steady state force before and after the step-release, the rate of both k_tr_ and Ph2 were independent of RLC phosphorylation. This confirms that the faster rate of force development and myosin head activation after RLC phosphorylation depends on the restretch and the mechanical load on the thick filament. This was also independent of the length of the applied step-release (**Fig. S6**).

In contrast to activating conditions, RLC phosphorylation had only a small effect on the RLC E-helix orientation during the relaxed state, which was largely independent of sarcomere length and passive force (**Fig. S7a**). Similarly, RLC phosphorylation had no effect on the ATPase activity of isolated relaxed rat ventricular myofibrils, suggesting that it does not affect the myosin head resting ATPase (**Fig. S7b**). This is in good agreement with previously published results, showing that the force-dependent activation of the myosin motors requires thin filament Ca^2+^-activation (11).

## Discussion

The results presented in the current study show that phosphorylation of the cardiac myosin motors in intact thick filaments is not a homogenous process but more likely a spatially tightly controlled mechanism that directly relates to its effects on sarcomere contractile function to increase myofilament calcium sensitivity and force development. Strikingly, our results show that myosin heads in the inner two-thirds of the thick filament, i.e. the region associated with cardiac myosin binding protein-C, are the primary target for phosphorylation by cMLCK and that RLC phosphorylation likely only leads to the activation of one of the myosin motors of the double headed myosin molecule. Functionally, RLC phosphorylation increases the force-dependent recruitment of the myosin motors from the functional OFF into the ON state, suggesting that phosphorylation of RLC increases the gain of the mechano-sensing mechanism of activation of the thick filament via increased signalling between the different thick filament zones as discussed in more detail below (45).

However, before discussing the implications of the current results for cardiac muscle function, we first consider the potential limitations of using exogenous cMLCK on isolated myofilaments or demembranated cardiac muscle cells as model systems. First, our experimental results are based on the catalytic subunit of recombinant cMLCK, which does not contain the N-terminal extension found in the full-length protein (18). Although the function of the N-terminal extension of cMLCK has not been characterized, it is conceivable that it might change its enzymatic activity, specificity or localization (46). Second, phosphorylation via exogenous cMLCK is calcium-dependent and was therefore performed in the presence of Blebbistatin to prevent myofilament contraction during the phosphorylation reaction (18). Blebbistatin stabilizes the OFF state of the myosin motors and thick filaments (47, 48), whereas in the intact heart the thick filaments continuously transition between the OFF and ON states (12). Third, isolation of cardiac myofibrils or demembranation of ventricular trabeculae could have led to a loss of soluble protein components that might affect RLC phosphorylation by cMLCK. However, given the similar spatial distribution of RLC phosphorylation in the sarcomeres of human myofibrils isolated under near-native conditions, we conclude that our experimental results are likely not an in-vitro artefact (Fig. S4).

An important question therefore is what controls the pre-dominant RLC phosphorylation in the thick filament C-zone? It seems likely that the intrinsic structure of the thick filament or the sarcomere itself regulates the spatial distribution of RLC phosphorylation. This is supported by the observation that the addition of exogenous kinase to isolated myofilaments is sufficient to phosphorylate myosin heads specifically in the C-zone, although the soluble kinase should be able to bind to all available binding sites in the sarcomere. It seems more likely that the different organization of the myosin motors on the surface of the thick filament backbone in the D- and C-zone might contribute to the zonal differences in phosphorylation by regulating the availability of the RLC N-terminal extension for cMLCK (35, 49). In agreement, both the steady state and kinetic measurements of RLC phosphorylation in cardiac myofibrils shown above suggest the presence of at least two populations of myosin heads that are phosphorylated with kinetics that differ by about an order of magnitude, which might represent heads in the D- and C-zone. Noteworthy, a previous study using isolated skeletal muscle myosin light chain kinase in mouse ventricular myofibrils showed a kinetically homogenous phosphorylation of RLC (19).

Another hypothesis is that cMyBP-C sequesters cMLCK to the C-zone via direct protein-protein interaction. Previous studies have shown that native cMyBP-C co-purifies from ventricular tissue with a Ca^2+^/calmodulin-dependent protein kinase, consistent with the Ca^2+^/calmodulin-dependent activity of cMLCK (50). However, other studies have shown that cMLCK is mostly localized to the cytosolic compartment of cardiomyocytes with only some minor striated appearance (19, 51, 52), although this does not exclude equilibrium binding of the kinase with moderate affinity. Additionally, interaction of cMLCK with other sarcomeric proteins such as titin or thin filament components might contribute to the spatial distribution of RLC phosphorylation by either localizing cMLCK to the C-zone or modulating its enzymatic activity.

The asymmetric structure of the myosin molecule OFF or IHM state suggests differential regulation of the activity of the two myosin heads (8, 30). The comparison of our fluorescence polarization studies in rat ventricular trabeculae with the recently published cryoEM structure of the human cardiac myosin IHM suggest that phosphorylation mainly affects the conformation of the RLC of the so called ‘blocked head’ of the myosin dimer (30) (**Fig. 3**). In contrast, the conformation of the ‘free head’ was not or less affected by phosphorylation. Although this seems contradictory with the original nomenclature based on electron microscopy of isolated invertebrate thick filaments, more recent cryoEM reconstructions of intact human cardiac thick filaments in the OFF-state show that only the ‘free heads’ interact with cMyBP-C (4, 8). An intriguing hypothesis is that cMyBP-C stabilizes the OFF state of the ‘free heads’ even after RLC phosphorylation, whereas the ‘blocked heads’ are more likely to leave the OFF state and become available for actin-binding. In agreement, genetic ablation of cMyBP-C has been shown to increase the functional effects of RLC phosphorylation in isolated mouse myocardium, likely by allowing the recruitment of both ‘free’ and ‘blocked’ heads for force generation (53). Similalrly, phosphorylation of a single RLC is sufficient to abolish the asymmetric inhibited IHM state of isolated dimeric smooth muscle myosin (54).

The potential implications of the above findings for the interpretation of the functional effects of RLC phosphorylation are summarized in **Figure 6**. The thick filament mechano-sensing hypothesis predicts that sentinel myosin heads in the D-zone at the tip of the thick filament attach early during calcium activation of myofilament, which creates filament strain that leads to the subsequent activation of the rest of the filament towards the M-line (**Fig. 6a**) (12, 45). The ends of the thick filaments lack cMyBP-C and likely have a different filament backbone configuration compared to the C-zone associated with the different titin repeats. Moreover, the thick filament backbone tapers off towards their tips at the A-I junction. Cumulatively, these factors likely lead to a de-stabilization of the myosin motor OFF configuration in the D-zone in comparison with the rest of the filament (**Fig. 6a, top**). It follows that calcium activation of the thin filaments will lead to myosin heads in the D-zone attaching to actin first, whereas the C-zone myosin heads remain in the OFF state (**Fig. 6a**, middle). Force generated by the D-zone motor is subsequently transmitted along the thick filament backbone to the C-zone which activates those motors and the rest of the thick filament (**Fig. 6a**, bottom).

**Figure 6.**
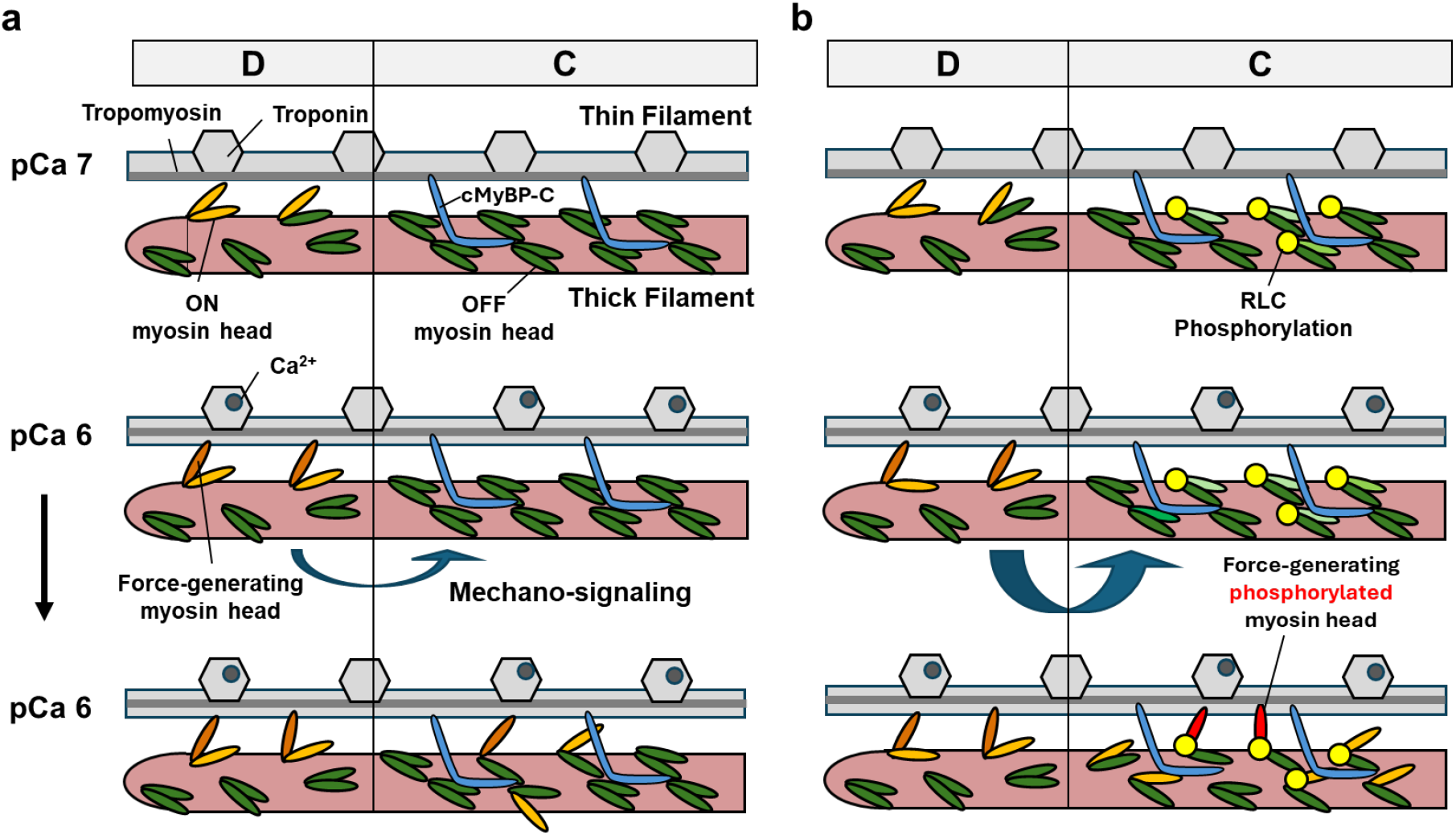
Hypothetical model for the effects of RLC phosphorylation on thick filament activation. (a) Activation of C-zone motors is triggered by force-generating myosin heads at the tip of the thick filaments (D-zone) via a mechano-signalling mechanism after calcium activation of the thin filaments. (b) RLC phosphorylation increases the mechano-signalling between the D- and C-zone, leading to an increased activation of myosin motors for the same level of calcium activation.

Although phosphorylation of RLC has no effect on the resting conformation of the myosin heads (**Fig. S7; Fig. 6b, top**), it significantly increases the sensitivity of the C-zone myosin motors to filament stress (**Fig. 6b, middle**), leading to a larger number of myosin motors being activated via mechano-signalling from the D-zone (**Fig. 6b, bottom**). Functionally, the consequence is a faster rate of and larger force-production for the same level of thin filament calcium activation, which is consistent with the increased calcium sensitivity of force and cross bridge cycling kinetics after RLC phosphorylation reported here and by others (22-24).

An interesting implication of this model is that the ‘free heads’ can likely be recruited for force-generation via additional mechanisms such as length-dependent-activation or phosphorylation of cMyBP-C (55, 56). In agreement, previous studies have shown that RLC phosphorylation and increase in sarcomere length have an additive and synergistic effect on the activation level of the myosin motors, calcium sensitivity and force production of isolated myocardium (24, 40). In this holistic view of regulation, the sarcomeric C-zone itself acts as a signal integrator that combines different inputs (e.g. filament stress, post-translational modifications and inter-filament spacing) to produce the appropriate output (i.e. force development).

Lastly, we consider the physiological relevance of the proposed mechanism for the modulation of sarcomere function during both health and disease states of the heart. The enzymatic activity of cMLCK is too slow to operate on a beat-to-beat basis and therefore it is more likely that RLC phosphorylation is part of a long-term adaptation that operates on the time scales of minutes to hours (19). Previous studies have reported average phosphorylation levels of about 0.4-0.5 mol P_i_ mol RLC^-1^ under basal conditions in rodent models. However, more recent studies showed an average of 0.25 mol P_i_ mol RLC^-1^ for human myocardial samples (19), whereas others have reported significantly lower levels for both (>0.1 mol P_i_ mol RLC^-1^ (57)). Nonetheless, taken together with our results, this suggests that under physiological conditions the mammalian heart primarily modifies the phosphorylation levels of myosin motors in the thick filament C-zone. Since myosin motors in the C-zone have been shown to produce the peak force of a cardiac twitch and their detachment kinetics are likely the primary determinant of mechanical relaxation, RLC phosphorylation allows direct control of both cardiac systolic force development and diastolic relaxation of the heart (12, 58).

Interestingly, the level of RLC phosphorylation is not altered during human heart failure (34), although others have reported changes in RLC phosphorylation in animal models of heart disease and heart failure (59). The current results suggest, however, that not only the total abundance of RLC phosphorylation is important but also its spatial distribution in the sarcomere and its dynamical changes.

## Methods

### Production of recombinant proteins

Recombinant proteins were prepared according to previously published protocols (33, 38, 60). Briefly, proteins were expressed from a modified pET6a vector fused to an N-terminal hexa-histidine tag and TEV protease site in BL21(DE3) cells at 18ºC for 18h using 0.1 mmol L^-1^ isopropyl β-D-thiogalactopyranoside (IPTG) in TB medium (Sigma Aldrich). Cells were harvested by centrifugation and lyzed using BugBuster Master Mix (Novagen) according to manufacturer’s instructions. Lysates were cleared by centrifugation at 10,000 g for 20 mins at 4ºC and filtration through a 0.22 μm syringe filter. The clear filtrate was applied to HisTrapFF columns (Cytiva) in binding buffer (composition in mmol L^-1^: 20 HEPES, 300 NaCl, 1 MgCl_2_, 1 DTT) and proteins eluted in buffer containing 250 mmol L^-1^ imidazole. The proteins were dialysed against buffer without imidazole and the hexa-histidine tag was removed by TEV protease cleavage. The solutions were applied to a second round of HisTrapFF purification to remove TEV protease, histidine-tag and uncleaved protein, and the flow-through collected. Proteins were further purified by ion-exchange chromatography on ResourceQ column according to manufacturer’s instructions.

The ADP biosensor was expressed from a pHis17_ParM_2 vector (kindly provided by Simone Kunzelmann, Crick Institute, United Kingdom) and expressed in BL21 Ai cells (Agilent Technologies) in 2×TY medium (Sigma-Aldrich) using 3 mg L^-1^ arabinose to induce protein expression at 30ºC a for 16h.The ParM was subsequently purified using HisTrapFF columns as described above. ParM was labelled with a four-times excess of tetramethylrhodamine-5-iodoacetamide (5-IATR) in 30 mmol L^-1^ Tris-HCl pH 7.5, 25 mmol L^-1^ KCl for 2h at room temperature. The labelled and purified ParM ADP biosensor was purified on a HiTrap Q column equilibrated in labelling buffer and proteins eluted using a linear gradient to 200 mmol L^-1^ KCl.

Double cysteine mutants of RLC were labelled with bifunctional sulforhodamine (BSR, Invitrogen) in 25 mmol L^-1^ HEPES, 1 mmol L^-1^ MgCl_2_ and the reaction monitored by HPLC (Agilent Technologies, 1100 system) and electron-spray ionization single quadrupole (ESI-Quad, Agilent Technologies) mass spectrometry. Labelled RLCs were purified on a MonoS column in labelling buffer using a linear gradient of 0-200 mmol L^-1^ KCl.

The catalitic subunit of cMLCK was prepared as described previously (18). Briefly, the fragment of human cMLCK [UniProtKB entry: Q32MK0] spanning amino acids 492-819 was expressed as an N-terminal fusion protein to Histdine-tag and TEV protease site in Spodoptera frugiperda 9 (Sf9) cells according to manufacturer’s instructions (BaculoDirectTM Baculovirus Expression System, Invitrogen). The fragment was purified by affinity chromatography on HIsTrapFF column (Invitrogen), followed by ion-exchange chromatography on CM-Sepharose (GE Healthcare) and gel filtration on a Superdex 75 HR 10/30 (GE Healthcare). The protein was concentrated to over 1 mg/ml and stored in 25 µl aliquots at -80^°^C for further use.

### Preparation of cardiac myofibrils

All animals were treated in accordance with the guidelines approved by the UK Animal Scientific procedures Act (1986) and European Union Directive 2010/63/EU. All procedures were performed according to Schedule 1 of the UK Animal Scientific Procedure Act, 1986, which do not require ethical approval. All procedures complied with the relevant ethical regulations and were carried out in accordance with the guidelines of the Animal Welfare and Ethical Review Body (AWERB, King’s College London).

Wistar rats (male, 200–250 g) were sacrificed by cervical dislocation without the use of anesthetics (Schedule 1 procedure in accordance with UK Animal Scientific Procedure Act, 1986) and demembranated right ventricular trabeculae were prepared as described previously (38).

Briefly, hearts were removed and rinsed free of blood in Krebs solution (composition in mmol L^−1^: 118 NaCl, 24.8 NaHCO_3_, 1.18 Na_2_HPO4, 1.18 MgSO_4_, 4.75 KCl, 2.54 CaCl_2_, 10 glucose, bubbled with 95% O_2_–5% CO_2_, pH 7.4 at 20 °C).

Cardiomyofibrils (CMFs) were prepared by homogenizing fresh ventricular tissue samples in myofibril buffer (composition in mmol L^−1^: 20 imidazole pH 7.4, 75 KCl, 2 MgCl_2_, 2 EDTA, 1 DTT, 1% (v/v) Triton X‐100, protease inhibitor cocktail (Roche), PhosStop cocktail (Roche)) followed by centrifugation at 5000 g for 5 min at 4°C. CMFs were washed and homogenized three more times in the same buffer without Triton X‐100.

### RLC phosphorylation assays

RLC and miniHMM were gel-filtered into cMLCK assay buffer (composition in mmol/L: 50 HEPES pH 7.0, 50 NaCl, 2 MgCl_2_, 1 CaCl_2_, 1 DTT) and protein concentrations adjusted to 25 µmol/L. cMLCK and calmodulin (CaM, kindly provided by Martin Rees, King’s College London) were added to final concentrations of 0.1 µmol L^-1^ and 0.3 µmol L^-1^, respectively, the reaction started by adding ATP to a final concentration of 1 mmol L^-1^. The reactions were incubated at 30°C, aliquots were quenched at the indicated time points with sample buffer and analysed by urea-glycerol-PAGE (4) or Phostag^TM^-SDS-PAGE. Bands were visualized by total protein staining with Coomassie. For the ADP biosensor assay, 20 μmol L^-1^ 5-IATR labelled ParM was added to the reaction mixture and the reaction monitored on a Clariostar plate reader (BMG LabTech) using appropriate excitation and emission filter settings.

Ventricular myofibrils (4 mg mL^-1^) were phosphorylated by exogenous cMLCK and CaM in activating solution (composition in mmol L^−1^: 25 Imidazole, 15 Na_2_CrP, 58.7 KPr, 6.3 MgCl_2_, 10 CaCl_2_, 10 K_2_EGTA, 1 DTT, pH 7.1) containing 25 μmol L^-1^ blebbistatin. The reactions were started by adding ATP to final concentration of 5 mmol L^−1^ and incubated at 30°C. Aliquots were quenched at the indicated time points with sample buffer and analysed by Phostag^TM^-SDS-PAGE and Westen-blot against RLC (primary antibody: rabbit monoclonal anti-myosin light chain 2,ABCAM, 1:dilution; secondary antibody: HRP-conjugated donkey anti-rabbit IgG, GE Healthcare, NA934V, 1:1000 dilution) as described previously (24). For the ADP biosensor assay, 10 μmol L^-1^ 5-IATR labelled ParM was added to the reaction mixture and the reaction monitored on a Clariostar plate reader (BMG LabTech) using appropriate excitation and emission filter settings.

### Super-resolution microscopy

Myofibrils were fixed with 4% (v/v) paraformaldehyde in PBS for 5 min at room temperature and subsequently washed three-times with PBS for 10 mins. Fixed myofibrils were incubated with primary antibodies (mouse anti-MHC clone A4.1025, Developmental Studies Hybridoma Bank - University of Iowa, 1:10 dilution; rabbit anti-phospho Ser15 RLC, 1:100, Affinity #AF8618) in labelling buffer (20 mmol L^-1^ Tris-HCl pH 7.5, 150 mmol L^-1^, 1 mg ml^-1^ bovine serum albumin) for 16 h at 4ºC in a humid chamber. Myofibrils were washed three-times for 10 mins in PBS and incubated with secondary antibodies (goat anti-rabbit-Alexa488, ABCAM #AB150077, 1:100 dilution; goat anti-mouse-Alexa594, ABCAM #AB150116, 1:100 dilution) in labelling buffer for 1 h at 25ºC. Myofibrils were washed in PBS, mounting medium (80% (v/v) glycerol, 40 mg mL^-1^ N-propyl gallat, 100 mmol L-1 Tris-HCl pH 8) was added and myofibrils sealed between microscope slides and cover glass using Tissue-TEK VIP embedding wax.

Myofibrils were imaged on a NIKON Spinning Disk confocal microscope with SoRA super-resolution module (CSU-W1 SoRA) according to manufacturer instructions using 60x objective.

### Preparation of cardiac trabeculae

Hearts were isolated from Wistar rats as described above for the preparation of cardiac myofibrils. Suitable trabeculae were dissected from the right ventricle in Krebs solution containing 25 mmol L−1 2,3-butanedione-monoxime, permeabilized in relaxing solution (see below) containing 1% (v/v) Triton X-100 for 30 min and stored in relaxing solution containing 50% (v/v) glycerol at −20 °C for experiments.

### Fluorescence polarization experiments

Trabeculae were mounted between a strain gauge force transducer (KRONEX, Oakland, California 94602, USA; model A-801, resonance frequency ∼2 kHz) and motor (Aurora Scientific, Dublin, D6WY006, Ireland; Model 312 C). BSR-cRLCs were exchanged into demembranated trabeculae by extraction in CDTA-rigor solution (composition in mmol L−1: 5 CDTA, 50 KCl, 40 Tris-HCl pH 8.4, 0.1% (v/v) Triton X-100) for 30 min followed by reconstitution with 40 μmol L^−1^ BSR-cRLC in relaxing solution (composition in mmol L^−1^: 25 Imidazole, 15 Na_2_Creatine phosphate (Na_2_CrP), 78.4 KPropionate (KPr), 5.65 Na_2_ATP, 6.8 MgCl_2_, 10 K_2_EGTA, 1 DTT, pH 7.1) for 1 h, replacing ∼50% of the endogenous cRLC.

Composition of experimental solutions and activation protocols were identical to those described previously for fluorescence polarization experiments. Polarized fluorescence intensities were measured as described previously for cardiac muscle fibres. Fluorescence emission from BSR-cRLCs in trabeculae were collected by a 0.25 N.A. objective using an excitation light beam in line with the emission path. The polarization of the excitation beam was switched at 1 kHz by a Pockels cell (Conoptics) between the parallel and perpendicular directions with respect to the muscle fibre long axis. The fluorescence emission was separated into parallel and perpendicular components by polarizing beam splitters, and its intensity measured by two photomultipliers, allowing determination of the order parameter <P2> that describes the dipole orientations in the trabeculae24. Force, muscle length and photomultiplier signals were constantly sampled at 10 kHz using dedicated programs written in LabView 2014 (National Instruments). Data were analysed using Microsoft Excel 2014 and GraphPad Prism 9.

The sarcomere length of trabeculae was adjusted to 2.2 μm by laser diffraction in relaxing solution prior to each activation. Activating solution contained (in mmol L^−1^): 25 Imidazole, 15 Na_2_CrP, 58.7 KPr, 5.65 Na2ATP, 6.3 MgCl_2_, 10 CaCl_2_, 10 K_2_EGTA, 1 DTT, pH 7.1. Each activation was preceded by a 2-min incubation in pre-activating solution (composition in mmol L^−1^: 25 Imidazole, 15 Na_2_CrP, 108.2 KPr, 5.65 Na_2_ATP, 6.3 MgCl_2_, 0.2 K_2_EGTA, 1 DTT, pH 7.1).

### FiberSim Modelling

The spatially explicit model FiberSim (41) was used to simulate force-pCa curves from different patterns of RLC phosphorylation. Myosin heads were assumed to cycle independently between the OFF, the ON and the force-generating state as shown in the kinetic scheme in Figure 4b. The three simulations are obtained by modulating the force-dependent transition between the OFF and the ON state to mimic different duty ratios between the two heads of a dimer and in different regions of the thick filament.

## Acknowledgements

The authors would like to thank Dr Xu Fu (University of Kentucky) for excellent support with the super-resolution imaging.

## Author Contributions

T.K. designed research; D.K., C.S. and T.K. performed research; T.K., K.S.C., C.S., D.K. and P.A. analyzed data; and T.K., K.S.C., C.S., D.K. and P.A. wrote the paper.

## Data Availability

The data supporting the findings of the study are available in the article and its Supplementary Information. All remaining raw data will be available from the corresponding author upon reasonable request.

## Code Availability

LabView programs used for data acquisition and analysis are available from the corresponding author upon reasonable request. The FiberSim code is freely available from https://campbell-muscle-lab.github.io/FiberSim/.

## References

1. A. M. Gordon, E. Homsher, M. Regnier, Regulation of contraction in striated muscle. Physiol Rev 80, 853–924 (2000).

2. M. Irving, Functional control of myosin motors in the cardiac cycle. Nat Rev Cardiol 10.1038/s41569-024-01063-5 (2024).

3. G. Piazzesi, M. Caremani, M. Linari, M. Reconditi, V. Lombardi, Thick Filament Mechano-Sensing in Skeletal and Cardiac Muscles: A Common Mechanism Able to Adapt the Energetic Cost of the Contraction to the Task. Front Physiol 9, 736 (2018).

4. J. L. Woodhead et al., Atomic model of a myosin filament in the relaxed state. Nature 436, 1195–1199 (2005).

5. L. Alamo et al., Three-dimensional reconstruction of tarantula myosin filaments suggests how phosphorylation may regulate myosin activity. J Mol Biol 384, 780–797 (2008).

6. M. E. Zoghbi, J. L. Woodhead, R. L. Moss, R. Craig, Three-dimensional structure of vertebrate cardiac muscle myosin filaments. Proc Natl Acad Sci U S A 105, 2386–2390 (2008).

7. H. A. Al-Khayat, R. W. Kensler, J. M. Squire, S. B. Marston, E. P. Morris, Atomic model of the human cardiac muscle myosin filament. Proc Natl Acad Sci U S A 110, 318–323 (2013).

8. D. Dutta, V. Nguyen, K. S. Campbell, R. Padron, R. Craig, Cryo-EM structure of the human cardiac myosin filament. bioRxiv 10.1101/2023.04.11.536274 (2023).

9. D. Tamborrini et al., Structure of the native myosin filament in the relaxed cardiac sarcomere. Nature 623, 863–871 (2023).

10. T. Kampourakis, M. Irving, The regulatory light chain mediates inactivation of myosin motors during active shortening of cardiac muscle. Nat Commun 12, 5272 (2021).

11. S. J. Park-Holohan et al., Stress-dependent activation of myosin in the heart requires thin filament activation and thick filament mechanosensing. Proc Natl Acad Sci U S A 118 (2021).

12. E. Brunello et al., Myosin filament-based regulation of the dynamics of contraction in heart muscle. Proc Natl Acad Sci U S A 117, 8177–8186 (2020).

13. J. Huang, J. M. Shelton, J. A. Richardson, K. E. Kamm, J. T. Stull, Myosin regulatory light chain phosphorylation attenuates cardiac hypertrophy. J Biol Chem 283, 19748–19756 (2008).

14. P. Ding et al., Cardiac myosin light chain kinase is necessary for myosin regulatory light chain phosphorylation and cardiac performance in vivo. J Biol Chem 285, 40819–40829 (2010).

15. S. B. Scruggs et al., Ablation of ventricular myosin regulatory light chain phosphorylation in mice causes cardiac dysfunction in situ and affects neighboring myofilament protein phosphorylation. J Biol Chem 284, 5097–5106 (2009).

16. S. Yadav et al., Therapeutic potential of AAV9-S15D-RLC gene delivery in humanized MYL2 mouse model of HCM. J Mol Med (Berl) 97, 1033–1047 (2019).

17. C. C. Yuan et al., Constitutive phosphorylation of cardiac myosin regulatory light chain prevents development of hypertrophic cardiomyopathy in mice. Proc Natl Acad Sci U S A 112, E4138–4146 (2015).

18. T. Kampourakis, M. Irving, Phosphorylation of myosin regulatory light chain controls myosin head conformation in cardiac muscle. J Mol Cell Cardiol 85, 199–206 (2015).

19. A. N. Chang et al., Constitutive phosphorylation of cardiac myosin regulatory light chain in vivo. J Biol Chem 290, 10703–10716 (2015).

20. E. Lee, Z. Liu, N. Nguyen, A. C. Nairn, A. N. Chang, Myosin light chain phosphatase catalytic subunit dephosphorylates cardiac myosin via mechanisms dependent and independent of the MYPT regulatory subunits. J Biol Chem 298, 102296 (2022).

21. A. N. Chang, K. E. Kamm, J. T. Stull, Role of myosin light chain phosphatase in cardiac physiology and pathophysiology. J Mol Cell Cardiol 101, 35–43 (2016).

22. B. A. Colson et al., Differential roles of regulatory light chain and myosin binding protein-C phosphorylations in the modulation of cardiac force development. J Physiol 588, 981–993 (2010).

23. M. C. Olsson, J. R. Patel, D. P. Fitzsimons, J. W. Walker, R. L. Moss, Basal myosin light chain phosphorylation is a determinant of Ca2+ sensitivity of force and activation dependence of the kinetics of myocardial force development. Am J Physiol Heart Circ Physiol 287, H2712–2718 (2004).

24. T. Kampourakis, Y. B. Sun, M. Irving, Myosin light chain phosphorylation enhances contraction of heart muscle via structural changes in both thick and thin filaments. Proc Natl Acad Sci U S A 10.1073/pnas.1602776113 (2016).

25. F. Sheikh et al., Mouse and computational models link Mlc2v dephosphorylation to altered myosin kinetics in early cardiac disease. J Clin Invest 122, 1209–1221 (2012).

26. J. S. Davis et al., A gradient of myosin regulatory light-chain phosphorylation across the ventricular wall supports cardiac torsion. Cold Spring Harb Symp Quant Biol 67, 345–352 (2002).

27. R. Craig, R. Padron, J. Kendrick-Jones, Structural changes accompanying phosphorylation of tarantula muscle myosin filaments. J Cell Biol 105, 1319–1327 (1987).

28. R. J. Levine, R. W. Kensler, Z. Yang, J. T. Stull, H. L. Sweeney, Myosin light chain phosphorylation affects the structure of rabbit skeletal muscle thick filaments. Biophys J 71, 898–907 (1996).

29. A. S. Khromov, A. V. Somlyo, A. P. Somlyo, Thiophosphorylation of myosin light chain increases rigor stiffness of rabbit smooth muscle. J Physiol 512 (Pt 2), 345–350 (1998).

30. A. Grinzato et al., Cryo-EM structure of the folded-back state of human beta-cardiac myosin. Nat Commun 14, 3166 (2023).

31. R. C. Venema, R. L. Raynor, T. A. Noland, Jr., J. F. Kuo, Role of protein kinase C in the phosphorylation of cardiac myosin light chain 2. Biochem J 294 (Pt 2), 401–406 (1993).

32. S. Kunzelmann, M. R. Webb, A fluorescent, reagentless biosensor for ADP based on tetramethylrhodamine-labeled ParM. ACS Chem Biol 5, 415–425 (2010).

33. S. Ponnam, T. Kampourakis, Microscale thermophoresis suggests a new model of regulation of cardiac myosin function via interaction with cardiac myosin-binding protein C. J Biol Chem 298, 101485 (2022).

34. B. C. W. Tanner et al., Sarcomere length affects Ca2+ sensitivity of contraction in ischemic but not non-ischemic myocardium. J Gen Physiol 155 (2023).

35. P. K. Luther et al., Understanding the organisation and role of myosin binding protein C in normal striated muscle by comparison with MyBP-C knockout cardiac muscle. J Mol Biol 384, 60–72 (2008).

36. H. L. Granzier et al., Deleting titin’s I-band/A-band junction reveals critical roles for titin in biomechanical sensing and cardiac function. Proc Natl Acad Sci U S A 111, 14589–14594 (2014).

37. J. Chandler et al., In situ FRET-based localization of the N terminus of myosin binding protein-C in heart muscle cells. Proc Natl Acad Sci U S A 120, e2222005120 (2023).

38. T. Kampourakis, Y. B. Sun, M. Irving, Orientation of the N- and C-terminal lobes of the Myosin regulatory light chain in cardiac muscle. Biophys J 108, 304–314 (2015).

39. U. A. van der Heide, S. C. Hopkins, Y. E. Goldman, A maximum entropy analysis of protein orientations using fluorescence polarization data from multiple probes. Biophys J 78, 2138–2150 (2000).

40. H. C. Pulcastro, P. O. Awinda, J. J. Breithaupt, B. C. Tanner, Effects of myosin light chain phosphorylation on length-dependent myosin kinetics in skinned rat myocardium. Arch Biochem Biophys 601, 56–68 (2016).

41. S. Kosta, D. Colli, Q. Ye, K. S. Campbell, FiberSim: A flexible open-source model of myofilament-level contraction. Biophys J 121, 175–182 (2022).

42. K. S. Campbell, P. M. L. Janssen, S. G. Campbell, Force-Dependent Recruitment from the Myosin Off State Contributes to Length-Dependent Activation. Biophys J 115, 543–553 (2018).

43. B. Brenner, E. Eisenberg, Rate of force generation in muscle: correlation with actomyosin ATPase activity in solution. Proc Natl Acad Sci U S A 83, 3542–3546 (1986).

44. R. E. Dale et al., Model-independent analysis of the orientation of fluorescent probes with restricted mobility in muscle fibers. Biophys J 76, 1606–1618 (1999).

45. M. Linari et al., Force generation by skeletal muscle is controlled by mechanosensing in myosin filaments. Nature 528, 276–279 (2015).

46. S. B. Scruggs, R. J. Solaro, The significance of regulatory light chain phosphorylation in cardiac physiology. Arch Biochem Biophys 510, 129–134 (2011).

47. T. Kampourakis, Z. Yan, M. Gautel, Y. B. Sun, M. Irving, Myosin binding protein-C activates thin filaments and inhibits thick filaments in heart muscle cells. Proc Natl Acad Sci U S A 111, 18763–18768 (2014).

48. T. Kampourakis, X. Zhang, Y. B. Sun, M. Irving, Omecamtiv Mercabil and Blebbistatin modulate cardiac contractility by perturbing the regulatory state of the myosin filament. J Physiol 10.1113/JP275050 (2017).

49. M. Gautel, O. Zuffardi, A. Freiburg, S. Labeit, Phosphorylation switches specific for the cardiac isoform of myosin binding protein-C: a modulator of cardiac contraction? EMBO J 14, 1952–1960 (1995).

50. H. C. Hartzell, D. B. Glass, Phosphorylation of purified cardiac muscle C-protein by purified cAMP-dependent and endogenous Ca2+-calmodulin-dependent protein kinases. J Biol Chem 259, 15587–15596 (1984).

51. O. Seguchi et al., A cardiac myosin light chain kinase regulates sarcomere assembly in the vertebrate heart. J Clin Invest 117, 2812–2824 (2007).

52. J. Y. Chan et al., Identification of cardiac-specific myosin light chain kinase. Circ Res 102, 571–580 (2008).

53. K. L. Turner, H. S. Morris, P. O. Awinda, D. P. Fitzsimons, B. C. W. Tanner, RLC phosphorylation amplifies Ca2+ sensitivity of force in myocardium from cMyBP-C knockout mice. J Gen Physiol 155 (2023).

54. B. A. Baumann et al., Phosphorylated smooth muscle heavy meromyosin shows an open conformation linked to activation. J Mol Biol 415, 274–287 (2012).

55. B. A. Colson et al., Protein kinase A-mediated phosphorylation of cMyBP-C increases proximity of myosin heads to actin in resting myocardium. Circ Res 103, 244–251 (2008).

56. X. Zhang et al., Distinct contributions of the thin and thick filaments to length-dependent activation in heart muscle. Elife 6 (2017).

57. Z. R. Gregorich et al., Comprehensive assessment of chamber-specific and transmural heterogeneity in myofilament protein phosphorylation by top-down mass spectrometry. J Mol Cell Cardiol 87, 102–112 (2015).

58. I. Morotti et al., An integrated picture of the structural pathways controlling the heart performance. Proc Natl Acad Sci U S A 121, e2410893121 (2024).

59. C. Toepfer et al., Myosin regulatory light chain (RLC) phosphorylation change as a modulator of cardiac muscle contraction in disease. J Biol Chem 288, 13446–13454 (2013).

60. T. Kampourakis, S. Ponnam, K. S. Campbell, A. Wellette-Hunsucker, D. Koch, Cardiac myosin binding protein-C phosphorylation as a function of multiple protein kinase and phosphatase activities. Nat Commun 15, 5111 (2024).

