## Supporting Information for "Spatial control of myosin regulatory light chain phosphorylation modulates cardiac thick filament mechano-sensing"

### Supporting Information Figures

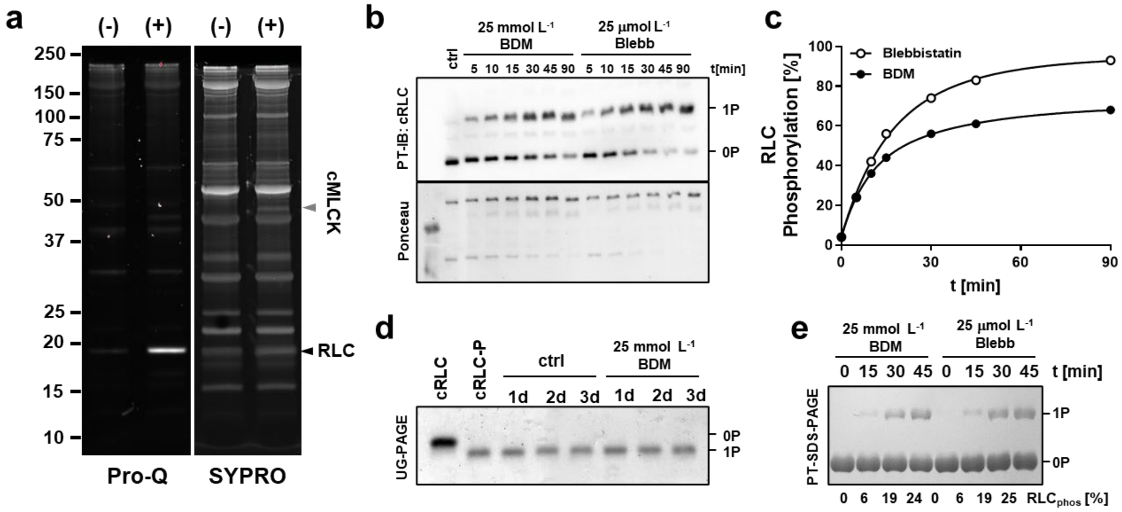


**Supporting Information Figure S1.** (a) Pro-Q Diamond and SYPRO Ruby staining of SDS-PAGE separating rat ventricular myofibrillar proteins before (-) and after cMLCK treatment (+). (b) Phostag™-Western-blot of the time-dependent phosphorylation of RLC in ventricular myofibrils in the presence of either butadiene, 2,3-monoxime (BDM) or blebbistatin (Blebb). (c) Time course of RLC phosphorylation shown in (b). (d) Isolated recombinant phosphorylated RLC was incubated in the absence (ctrl) or in the presence of 25 mmol $L^{-1}$ BDM for up to three days at 25ºC, and the RLC phosphorylation level determined by urea-glycerol PAGE (UG-PAGE). (e) Effect of BDM or Blebb on the phosphorylation of isolated RLC by cMLCK analyzed by Phostag™-SDS-PAGE. The relative amount of phosphorylated RLC is indicated below the gel.

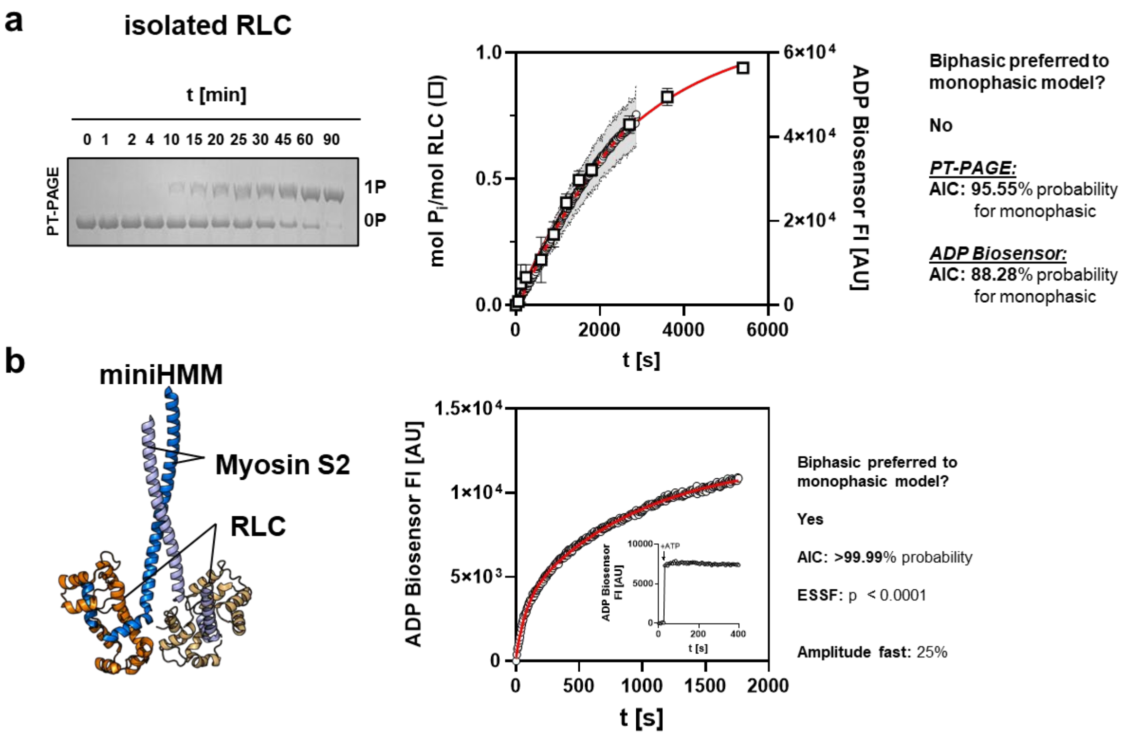


**Supporting Information Figure S2.** (a) Time-dependent phosphorylation of isolated RLC by cMLCK analyzed by Phostag$^{TM}$-SDS-PAGE (white squares) and ADP-biosensor assay (white circles). Red continuous line denotes fit to a mono-exponential function. (b) Time-dependent phosphorylation of isolated miniHMM by cMLCK analyzed by ADP-biosensor assay (white circles). Red continuous line denotes fit to a bi-exponential function. Inset shows control experiments in the absence of miniHMM with a stable baseline.

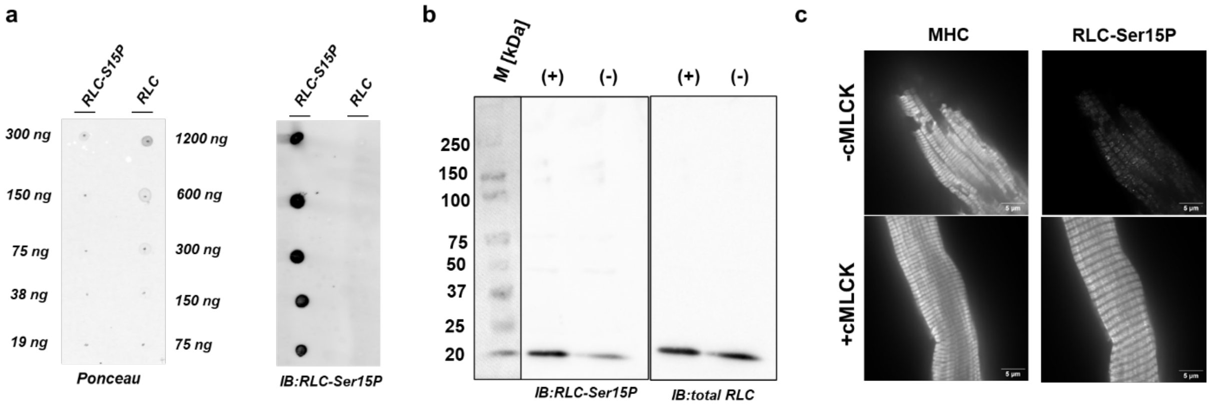


**Supporting Information Figure S3. Antibody validation.** (a) Dot-blot of serine 15 phosphorylated (RLC-Ser15P) and unphosphorylated recombinant rat ventricular RLC using the anti-RLC-Ser15P antibody. Loaded mounts of recombinant protein are indicated accordingly. (b) Western-blot of myofibrils samples before (-) and after (+) cMLCK treatment using the anti-RLC-Ser15P antibody (left) and total RLC antibody (right). (c) Confocal images of myofibrils before (-MLCK) and after cMLCK (+cMLCK) treatment stained against myosin heavy chain (MHC) and serine 15 phosphorylated RLC.

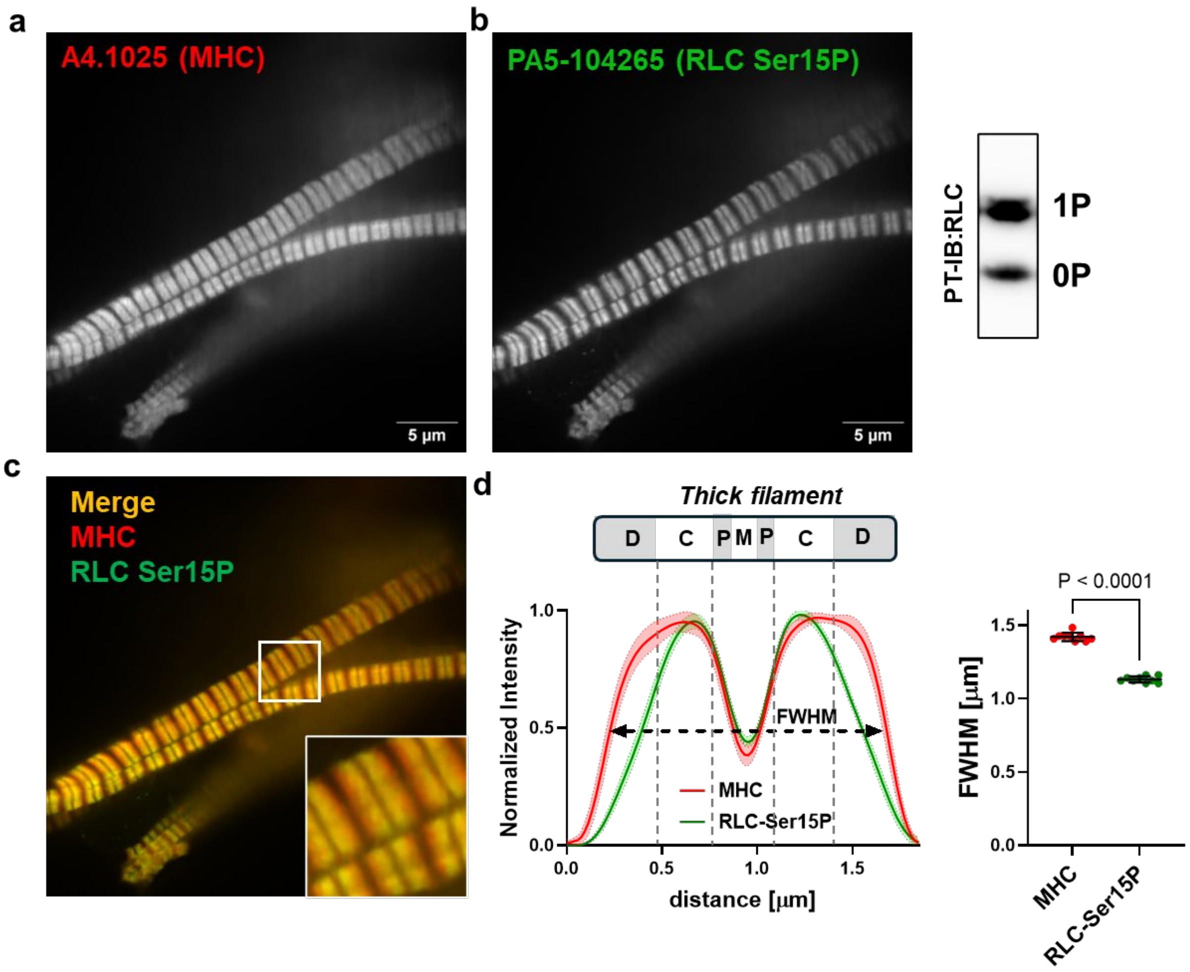


**Supporting Information Figure S4. Spatial RLC phosphorylation distribution in human ventricular myofibrils under native conditions.** (a) Stain for myosin heavy chain (MHC, red). (b) Left: Stain for Serine 15 phosphorylated RLC (RLC Ser15P, green). Right: Phostag$^{TM}$-Western-blot analysis of RLC phosphorylation levels in human ventricular myofibrils. (c) Merge of (a) and (b). (d) Left: The normalized averaged intensity profiles over n=9 sarcomeres. Continuous lines denote average profiles and shaded areas indicated 95% confidence intervals. The structure of the thick filament is shown to scale above the plot and the different filament zones are labelled accordingly. Right: Full width half maximum (FWHM) of the MHC and RLC-Ser15P distribution. Statistical significance of differences was assessed with an unpaired, two-tailed student's t-test (n=9 for MHC and n=9 for RLC-Ser15P).

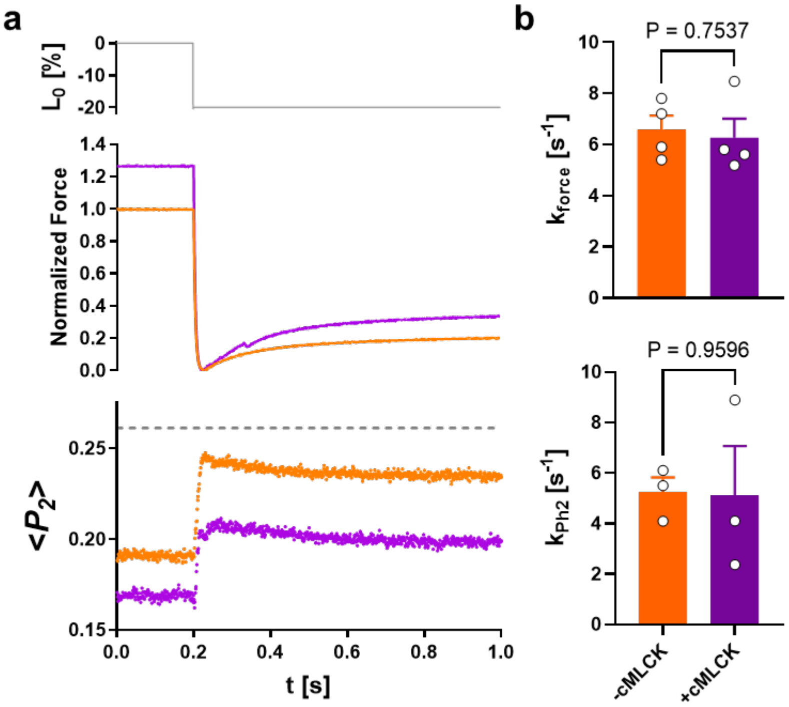


**Supporting Information Figure S5. Changes in the orientation of the cRLC E-helix probe in ventricular trabeculae in response to step shortening before and after cMLCK treatment.** (a) Representative traces of muscle length (top), force (middle) and $<P_2>$ before (orange) and after RLC phosphorylation (purple). (b) Summary of $k_{tr}$ (n=4 independent trabeculae preparations) and Ph2 rates (n=3 independent trabeculae preparations). Statistical significance of differences between values were assessed with a paired, two-tailed student's t-test.

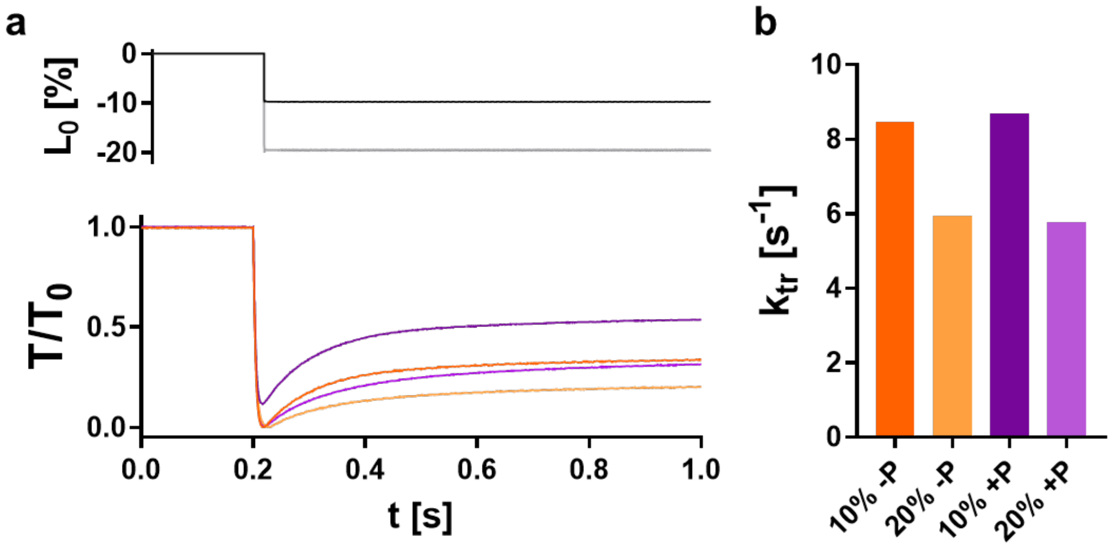


**Supporting Information Figure S6.** Rate of force re-development of ventricular trabeculae after 10% and 20% shortening steps before (orange) and after RLC phosphorylation (purple). Representative traces are shown in (a) and data summarized in (b) for n=1 preparation.

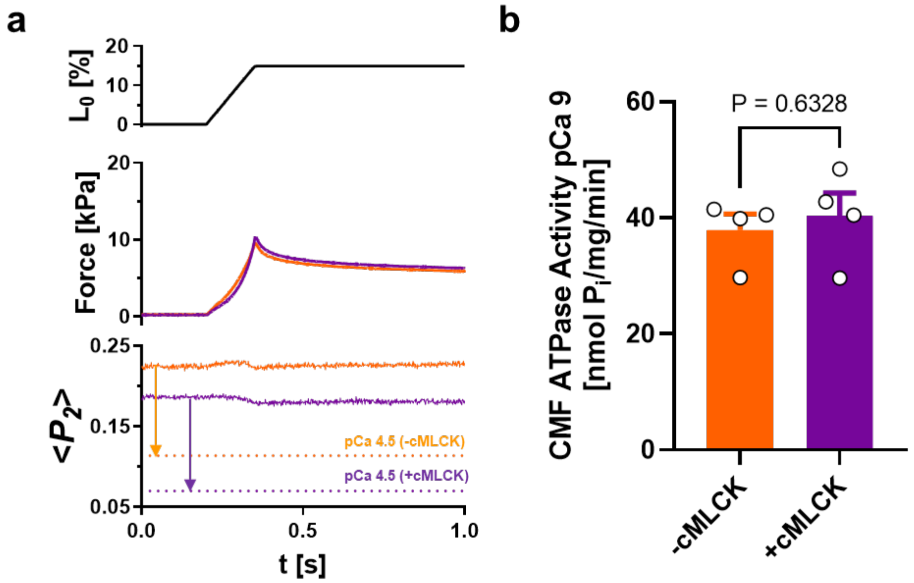


**Supporting Information Figure S7.** Effect of RLC phosphorylation on (a) relaxed myosin head orientation in ventricular trabeculae 9 (n=1) and (b) ATPase activity of isolated myofibrils (n=4 independent preparations) at pCa 9. Statistical significance of differences between values were assessed with a paired, two-tailed students' t-test.
